# Phylogenomics and macroevolution of a florally diverse Neotropical plant clade (Hillieae)

**DOI:** 10.1101/2025.03.29.645733

**Authors:** Laymon D Ball, Ana M Bedoya, Charlotte M Taylor, Laura P Lagomarsino

## Abstract

Hillieae is a group of ∼30 florally diverse, Neotropical epiphyte species. Species richness peaks in southern Central America and taxa display bat, hawkmoth, or hummingbird pollination syndromes. A phylogenetic framework is needed to understand floral and biogeographic evolution. We used target enrichment data to infer a species tree and a Bayesian time-calibrated tree including ∼83% of the species in the group. We inferred ancestral biogeography and pollination syndromes, described species realized bioclimatic niches via a principal component analysis, and estimated significant niche shifts using Ornstein-Uhlenbeck models to understand how different abiotic and biotic variables have shaped Hillieae evolution. We estimated that Hillieae originated in southern Central America 19 Ma and that hawkmoth pollination is the ancestral character state. Multiple independent shifts in pollination syndrome, biogeographic distribution, and realized bioclimatic niche have occurred, though bioclimatic niche is largely conserved. Using generalized linear models, we identify two interactions— between species’ biogeographic ranges and pollination syndromes, and between phylogenetic covariance and pollination syndromes— that additively affect the degree of bioclimatic niche overlap between species. Regional variation in pollination syndrome diversity and patterns of species bioclimatic niche overlap indicate a link between biogeography and species ecology in driving Hillieae diversification and syndrome evolution.

## Main

Altogether, the Neotropics comprise an estimated 118,000 flowering plant species, amounting to 40% of the world’s known flora (Raven et al. 2020). Macroevolutionary studies to date have consistently underscored the important roles of multiple abiotic and biotic factors in generating this biodiversity, including mountain uplift, barriers to dispersal, evolution of growth forms (especially epiphytism), mutualistic interactions between plants and pollinators, and antagonistic interactions such as herbivory and nectar-robbing (Givnish et al. 2014, 2015; Lagomarsino et al. 2016; Maron et al. 2019; Dellinger et al. 2024). The contribution of each factor to diversification may vary with environmental context (Vargas et al. 2020). For example, in lineages inhabiting highly mountainous regions like the Andes, abiotic factors may play a more significant role. Mountains can act as physical barriers to dispersal, fostering geographic speciation. Surface uplift can also alter environmental variables such as climate, drainage pathways, and soil conditions, generating new environmental niches that can foster local adaptation and speciation (Hoorn et al. 2010). In other regions with less topography, specialized ecological interactions may have played a larger role in structuring evolutionary relationships than the abiotic environment (Verboom et al. 2015).

Rubiaceae is one of the largest and most diverse plant families, comprising ∼13,500 species spread across ∼620 genera and three subfamilies (POWO, 2025). The family has a cosmopolitan distribution but is mainly tropical. It includes several ecologically and economically important plants such as coffee, quinine, and gardenias. Within the subfamily Cinchonoideae, the Hillieae tribe is a small but morphologically and ecologically diverse group of plants that has received relatively little attention in the scientific literature (Knudsen and Tollsten 1993; Taylor 1994; Sazima et al. 1999; D’hondt et al. 2004; Wolff 2006; Ramírez and Briceño 2021). This group comprises ∼30 mostly shrubby, epiphytic species distributed among the genera *Balmea* (1 species), *Cosmibuena* (4 species), and *Hillia* (∼25 species). Despite its relatively low species richness, the group exhibits remarkable floral diversity. This suggests that pollinator-mediated selection has likely played an important role in evolution, resulting in species displaying bat, hawkmoth, and hummingbird pollination syndromes, with observed intermediates. Most species lack field observations of pollinators, but four species— *C. valerii* (hawkmoth), *H. illustris* (bat), *H. parasitica* (hawkmoth), and *H. triflora* var. *triflora* (hummingbird)— have documented pollinator visits and their observed pollinators match expectations based on pollination syndromes (Knudsen and Tollsten 1993; Sazima et al. 1999; Wolff 2006; Ramírez and Briceño 2021; personal observation).

A distinguishing trait of Hillieae among other Rubiaceae is the presence of tiny, flattened, trichome-tufted (i.e., comose) seeds that develop within woody capsular fruits and are wind-dispersed. This seed morphology, which is unique to the genus *Hillia*, may be an adaptation that allows seeds to stay in the air for longer and travel farther, even relative to other Hillieae (Taylor 1994). Hillieae are broadly distributed throughout most of the Neotropics and most species occur in Southern Central America and the Northern Andes of South America (Taylor 1994). Species mainly occur in moist or wet low-to-mid elevation forests; however, some species can occur at relatively high elevations and in drier habitats.

Despite its marked floral and biogeographic diversity, phylogenetic relationships within Hillieae have been understudied. While a few species have been included in molecular phylogenetic studies, those have focused on evolutionary relationships and patterns across Rubiaceae broadly (Robbrecht and Manen 2006; Manns and Bremer 2010; Manns et al. 2012; Paudyal et al. 2014). These studies agree on the placement of Hillieae within subfamily Cinchonoideae, monophyly of the three Hillieae subgenera, and the placement of *Balmea* and *Hillia* as sister genera. Hillieae has also been the focus of multiple morphological cladistic analyses (Taylor 1994; D’hondt et al. 2004). However, morphological characters can mislead phylogenetic analyses when convergently evolved traits are included in data matrices, as may be the case in Hillieae. Consequently, our knowledge of relationships within Hillieae has remained uncertain.

Given its high floral diversity and occurrence in various habitats across the Neotropics, both abiotic and biotic factors have likely strongly influenced Hillieae diversification. To understand evolutionary relationships in the tribe and assess how biogeography and plant-pollinator interactions have shaped its evolution, we infer the first deeply sampled, time-calibrated, molecular phylogeny for Hillieae. We reconstruct historical biogeography, describe species’ bioclimatic niche shifts, estimate pairwise niche overlap between species, and reconstruct pollination syndrome evolution. Additionally, we apply generalized linear models (GLMs) to assess how different variables— including geographic co-occurrence, shared pollination syndrome, and phylogenetic relatedness— both independently and interactively affect the degree of niche overlap among pairs of species. Our findings suggest that although dispersal, shifts into different environmental niches, and changes in pollination syndromes have facilitated diversification, the weight of these mechanisms seems to vary geographically.

## Materials and Methods

### Gene Tree Inference

Taxon sampling, DNA extraction protocol, sequencing and locus assembly follow Ball et al. (Ball et al. 2023). Briefly, we performed target enrichment sequencing using a custom, Rubiaceae-specific probe set (Rubiaceae2270x) designed to target 2270 exonic loci (Ball et al. 2023), which were assembled via HybPiper 2 (Johnson et al. 2016). We included all Hillieae taxa with assembled contigs for at least 10% of loci, as well as all Hamelieae (Cinchonoideae) taxa and two species from subfamily Rubioideae (*Palicourea attenuata* and *Psychotria panamensis*). This sampling represents 25 Hillieae species, or ∼83% of all species in the tribe (Supplementary Table S1).

The HybPiper 2 “paralog_retriever” command was used to extract all assembled contigs for all loci, and we identified as putative paralogs those loci for which two or more contigs were assembled across samples. Sets of both putatively single-copy and paralogous loci were aligned using macse (version 2.07; Ranwez et al. 2018) under default settings, and then alignments were visually inspected in Geneious Prime (version 2021.1.1). Putatively paralogous loci were then removed from the dataset and processed separately (see next paragraph). For the remaining putatively single-copy loci, erroneous sequences from alignments (i.e., short segments of DNA from a given species that appear nearly random relative to the rest of the alignment) were removed with TAPER (version 1.1.1; Zhang et al. 2021), columns with more than 80% missing data were removed with the pxcslq function in phyx (version 1.1; Brown et al. 2017), and sequences shorter than 20% of the total alignment length were removed using trimAl (version 1.2; Capella-Gutiérrez et al. 2009). Alignments with less than 25 taxa (>60% missing taxa) were removed from downstream analyses.

Putatively paralogous loci were processed using a modified version of the Monophyletic Outgroups (MO) pipeline of Morales-Briones et al. (Morales-Briones et al. 2022; https://github.com/ambed0ya/Palicourea). First, columns with more than 80% missing data were removed with the pxcslq function in phyx, and spurious sequences were removed with trimAl. Preliminary gene trees were inferred using the program RAxML-NG (Kozlov et al. 2019) under the GTR+G nucleotide evolution model. Ortholog trees were then inferred by pruning trees with monophyletic and single-copy outgroup taxa. During orthology inference, we chose to only retain gene trees with at least 25 (60%) taxa. Sequence matrices were then generated from the MO-pruned gene trees (i.e., the cleaned original alignments were realigned to only include taxa remaining in the pruned gene tree). These alignments were again visually inspected in Geneious, and erroneous sequences were removed with TAPER. Columns with more than 80% missing data were removed with the pxcslq function in phyx, and spurious sequences were removed with trimAl. RAxML-NG was used to infer the final gene trees using the GTR+G model. We calculated summary statistics for all final processed alignments with AMAS (version 1.1.0; Borowiec 2016; Supplementary Table S3). RAxML-NG was used to infer gene trees using the GTR+G model.

### Phylogenetic Inference and Time Calibration

Phylogenetic trees were inferred using multiple approaches. We performed species tree inference analyses using the weighted-ASTRAL (i.e., wASTRAL) algorithm (Zhang and Mirarab 2022), a two-step coalescence aware approach. wASTRAL is a version of ASTRAL (Zhang et al. 2018) with weighting schemes to reduce the impact of quartets with low support and long terminal branches. All final gene trees (i.e., single-copy and inferred orthologs) were used as input. Phylogenetic relationships were also inferred in a concatenation analysis using RAxML-NG. In addition to the local posterior probability (LPP) values estimated with wASTRAL, Quartet Sampling (Pease et al. 2018) was used to calculate support for the wASTRAL species tree topology and distinguish lack of support from gene-tree conflict (--align concatenated.phylip --reps 200 --threads 4 --lnlike 2). For the concatenation analysis, all final processed alignments were concatenated into a single matrix, and a tree was inferred under the GTR+G model of nucleotide evolution. Branch support was estimated via the calculation of Felsenstein’s bootstrap proportions (FBP).

We estimated divergence times for our inferred species tree with RevBayes (version 1.2.1; Höhna et al. 2016). We used a secondary calibration on the crown node of Hillieae, derived from previous study of tribe Cinchonoideae where four fossil calibration constraints were specified to infer divergence times (Manns et al. 2012). We acknowledge that divergence times estimates are subject to many biases, and additional caveats come with using secondary calibrations (see Schenk 2016). Therefore, these estimates should be taken cautiously and viewed as a means of exploring plausible evolutionary scenarios.

To subset our loci and make the calibration more computationally feasible, we used the program genesortR (Mongiardino Koch 2021). GenesortR ranks loci by first calculating seven locus properties (pairwise patristic distance, compositional heterogeneity, level of saturation, root-to-tip variance, Robinson-Foulds similarity to a target topology [here, we use the wASTRAL species tree], average bootstrap support, and the proportion of variable sites) and then it uses a principal component analysis to identify an axis where proxies for phylogenetic signal increase and sources of bias decrease. We limited the genesortR search to 15 loci that included all 37 ingroup taxa. Next, we used the *chronos* function from the *ape* R package (version 5.7.1; Paradis and Schliep 2019) to generate an ultrametric starting tree compatible with our secondary calibration. We used the wASTRAL tree as input after pruning it to only include one individual per species and exclude all outgroups. For each species, we retained the individual with the most assembled loci, except for *H. macrophylla* for which we kept two individuals that were polyphyletic and sampled from geographically distant regions (i.e., Costa Rica and Ecuador). We applied a relaxed clock model and normal distribution with a minimum and maximum root age of 13.6 and 28.8 Ma respectively. In the RevBayes analysis, we specified a partitioned GTR+G substitution model, an exponential relaxed clock model, and a birth-death tree prior. We ran two independent chains for 300,000 generations and discarded 25% of posterior trees. A maximum clade credibility tree was generated from the remaining posterior distribution of trees. Convergence was assessed by visually inspecting trace files in the program TRACER (v.1.7.1; (Rambaut et al. 2018) and checking that all parameters had good mixing (effective sample size >200).

### Biogeographic Modeling in Hillieae

We used BioGeoBEARS (version 1.1.3; Matzke 2013) for biogeographic reconstruction under the Dispersal Extinction Cladogenesis (DEC; (Ree and Smith 2008), BayArea-like (Landis et al. 2013), and DIVA-like (Ronquist 1997) models using our time-calibrated phylogeny. To define species’ ranges, we retrieved occurrence data for all Hillieae species with accessible locality information from GBIF (https://doi.org/10.15468/dl.wjanz2), focusing on specimens from the following herbaria: COL, CR, F, INB, MO, NY, and US. The *clean_coordinates* function from the *CoordinateCleaner* R package (version 3.0.1; Zizka et al. 2019) under default settings was used to filter out erroneous or invalid records. Occurrence points were also compared with published species distributions (Taylor 1989, 1992, 1994; Taylor and Gereau 2010). In addition to the coordinates retrieved from GBIF, we also estimated latitude and longitude for two additional specimens of *Hillia bonoi* and two additional specimens of *H. macbridei*, which had particularly few total georeferenced records on GBIF, using locality information from the specimen labels. Based on our inferred species tree topology (see ‘Phylogenetic Inference and Time Calibration’ results below), the Central American and Andean collections of *H. macrophylla* were treated as separate operational taxonomic units.

We defined the bioregions included in the analysis as: Northern Central America, Southern Central America, the Andes, and Amazonia (Fig. 2a). These regions have known importance for Neotropical flora and were selected based on Hillieae distribution patterns in georeferenced collections. While multiple distantly related species also occur in the Caribbean *(H. parasitica* and *H. tetrandra*) and Atlantic Forest (*H. illustris*, *H. parasitica*, and *H. ulei*), we excluded these areas from the analysis to limit parameters, since they hold no endemic species and are areas occupied by Hillieae taxa with broad distribution ranges. For each model, we set the maximum number of areas that an ancestor could occupy to three, which is the greatest number of areas that any extant Hillieae species occupies. Dispersal rates between regions were scaled based on the shortest distances between their edges, which were calculated in QGIS (version QGIS 3.14.15) using the expression length (shortest_line($geometry, geometry(get_feature()))) in the field calculator. The distance between adjacent regions was set to 1 km. Disjunct ranges were constrained by manually modifying the list of possible states. The three models were compared using the AIC.

### Realized Bioclimatic Niche Evolution

To describe species’ realized bioclimatic niches, raster layers for all 19 bioclimatic variables and elevation were first downloaded from the WorldClim (https://www.worldclim.org/data/worldclim21.html) online historical climate database at a resolution of 2.5 arc-minutes. The 20 raster layers were clipped to the extent of Hillieae’s distribution using the GDAL “clip raster by extent” function in QGIS (version 3.22.4). After clipping the rasters, we used the *raster.cor.plot* function of the *ENMTools* R package (version 1.1.1; Warren et al. 2021) and the *removeCollinearity* function of the *virtualspecies* package (version 1.6; Leroy et al. 2016) to identify and filter out strongly correlated variables. This left a remainder of five bioclimatic variables for downstream analyses: annual precipitation, annual mean temperature, elevation, precipitation seasonality, and temperature seasonality.

To summarize and visually inspect the available environmental conditions based on the five filtered variables, we performed a principal component analysis (PCA) on the 455,929 land pixels covering Hillieae’s distribution (excluding the Caribbean and Atlantic forest) (Fig. 2c), following Alexandre (Alexandre et al. 2017). We considered the first two axes of the PCA, which explain most of the variation in the data. For each species with at least five occurrence records, we used the geographic coordinates retrieved in the previous section to identify the land pixels in which the taxon occurs (using the *cellFromXY* function of the *rts* package [version 1.1.14]) and then calculated the average score of these pixels on both dimensions of the PCA. These average scores were used as a proxy for the realized bioclimatic niche of the respective species. Additionally, we tested for significant differences among individuals with bat, hawkmoth, and hummingbird pollination syndromes on each climatic PCA axis using two approaches: (1) ANOVA, incorporating all species from the climatic PCA and including species assignment as a random effect; and (2) a phylogenetically informed analysis using the *MCMCglmm* R package (version 2.36; Hadfield 2010), including only species present in both the climatic PCA and the phylogeny. For the *MCMCglmm* analysis, we used our time-calibrated phylogeny as input, included species assignment as a random effect, and kept all other settings at default values. An R script used to perform the analyses is available at https://github.com/laymonb/Hillieae_macroevolution.

To estimate niche overlap across taxa, we estimated Schoener’s D index using the *ecospat.niche.equivalency.test* function from the *ecospat* R package (version 4.0.0; Di Cola et al. 2017). We included all species with at least five occurrence records that were sampled in the phylogeny. Using generalized linear models (GLMs), we tested whether sharing the same pollination syndrome, occurring in the same biogeographic region, being closely related, or an interaction between these variables could explain patterns of niche overlap. Phylogenetic covariance, calculated using the *vcv.phylo* function from the *ape* R package, was used as a measure of relatedness. In total, we tested 17 models (Supplementary Table S8) using the *glmmTMB* R package (version 1.1.10; Brooks et al. 2017). For each model, we applied a Beta family distribution and a logit link. We used BIC to compare the full set of models; for models with similar BIC values, we used the R DHARMa package (version 0.4.7) to compute additional diagnostic statistics and evaluate model fit.

Finally, we used the *estimate_shift_configuration* function from the *l1ou* R package (Khabbazian et al. 2016) to detect evolutionary shifts in bioclimatic niche optimum. Briefly, the function fits multi-optima Ornstein-Uhlenbeck (OU) models using the lasso method (Tibshirani et al. 2012) to detect shifts in phenotypic optima across a phylogeny, without quantifying if optimal values are reached. For every species included in the phylogeny (including species with few occurrences— *H. bonoi*, *H. foldatsii*, and *H. macbridei*), we calculated the median of the log10-transformed values for each of the five filtered bioclimatic variables used in the climatic PCA, and these median values were used as input for the analysis. We raised the “alpha.upper” parameter to eight to improve model convergence, and we used the phylogenetic-aware Bayesian information criterion (pBIC; Khabbazian et al. 2016) for model comparison. All other parameters were kept at default settings. Bootstrap support for shift positions was calculated using the *l1ou_bootstrap_support* function with 1000 replicates.

### Pollination Syndrome Evolution

We performed an ancestral state reconstruction of pollination syndromes to better understand how floral morphology has evolved through time. Floral diversity is a hallmark of Hillieae, and this variation reflects pollination mode diversity within the tribe. Multiple species (including *H. illustris*, *H. triflora*, *H. parasitica*, and *H. wurdackii*) have been included in pollination studies where reported visits match expectations based on the classic definitions of the bat, hawkmoth, and hummingbird pollination syndromes (Faegri and Pijl 1979; Knudsen and Tollsten 1993; Sazima et al. 1999; Wolff 2006; Ramírez and Briceño 2021). This suggests that flower color and shape are good proxies for pollination mode in Hillieae, and these traits were used to assign Hillieae species to syndromes. Species with red corollas (regardless of shape) were assigned the hummingbird syndrome, species with green and funnelform corollas were assigned the bat syndrome, and species with white and salverform corollas were assigned the hawkmoth syndrome. For the analysis, we first estimated the optimal model of character evolution using different functions from the *phytools* R package. Four models were run using equal rates (ER): a continuous-time Markov (Mk) model using the *fitMk* function; two hidden rates models with one and two hidden states using the *fitHRM* function; and an Mk model with edge rates assumed to have been randomly sampled from a Γ distribution using the *fitgammaMk* function. The same four models were also run using symmetrical rates (SYM) and all rates different (ARD), for a total of 12 models, which were compared using the AIC. Next, we inferred ancestral states for pollination syndrome in RevBayes under the best-fit model (i.e., continuous-time Mk model with equal rates). We ran two Markov chains for 10,000 generations each, sampling a stochastic character map every generation. We then summarized the 20,000 stochastic character maps generated from the two Markov chains using the *describe.simmap* function from the *phytools* package.

To better understand whether multiple independent origins of hummingbird and bat pollination syndromes could reflect hemiplasy (i.e., apparent trait convergence driven by gene trees underlying the trait of interest and discordant with the species tree [Avise and Robinson 2008; Guerrero and Hahn 2018]) rather than true convergence, we used a custom R script (https://github.com/laymonb/Hillieae_macroevolution) to calculate the number of gene trees from the total 1375 trees in our dataset in which species inferred to be distantly related in the species tree but that share pollination syndromes were inferred to be monophyletic. Gene trees were rooted on the most distant outgroup available prior to the analysis using the pxrr function from phyx.

## Results

### Gene Tree Inference

HybPiper assembly statistics for the 42 samples included in the present study are shown in Supplementary Table S2. 681 loci were recovered as putatively single-copy and 1578 exonic regions (loci) identified as putative paralogs were processed via the MO pipeline, resulting in an additional 801 inferred orthologs, for a total of 1482 loci. After alignment, cleaning, and filtering for loci with at least 25 taxa, we were left with a total of 1375 loci for phylogenetic inference. Alignments included 25–42 taxa (x^̅^=36 taxa) of 99–4791 bp length (x^̅^=328 bp), 0–47% missing data (x^̅^=3.7%), and 0–243 parsimony informative sites (x^̅^=40 sites).

### Phylogenetic Inference and Time Calibration

Relationships between Hillieae genera and *Hillia* subgenera in the wASTRAL and RAxML-NG topologies were fully congruent. However, the placement of some individuals within clades varied slightly (Figs. 1 and S1). Specifically, the trees were discordant in the monophyly of *Cosmibuena grandiflora* and *Hillia maxonii*, and in the placement of several taxa within *Hillia* subg. *Hillia*. In both trees, *Cosmibuena* and *Hillia* were recovered as monophyletic, and *Balmea* and *Hillia* as sister taxa. Bootstrap support in the RAXML-NG topology was 100 at all branches, except for four in the *Hillia* subg. *Hillia* clade in which bootstrap support ranged from 62–100 (x^̅^=0.98; Supplementary Fig. S1). LPP support in the wASTRAL species tree ranged from 0.67–1 (x^̅^=0.99) across all branches (Fig. 1). Five branches had an LPP less than 0.98 and all but one of these were concentrated in the *Hillia* subg. *Hillia* clade.

**Figure 1.**
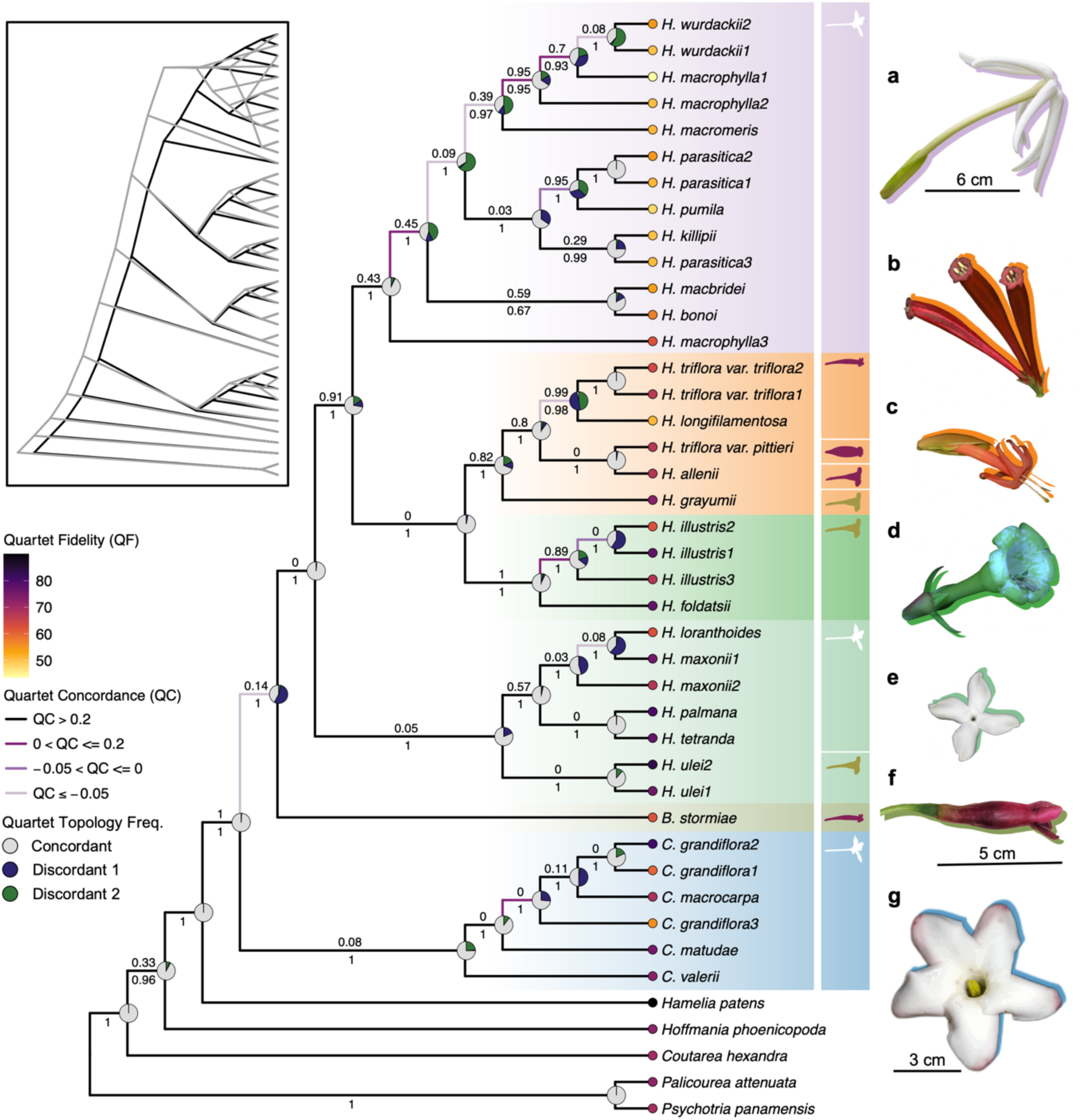
Phylogenetic tree inferred using wASTRAL with QuartetSampling (QS) support measures. Branch color indicates Quartet Concordance (QC) score, an entropy-like measure quantifying how often the concordant quartet was inferred over both discordant ones. Pie charts at nodes show the frequencies of QS replicates supporting concordant (white) and discordant (green and blue) topologies. The Quartet Differential (QD) score, which measures the skewness in the frequencies of the two discordant quartet topologies, is plotted above branches, except for instances where all quartets are concordant. wASTRAL LPP scores are plotted below branches. Colors at the tips indicate the Quartet Fidelity (QF) score, which reports the frequency of a taxon’s inclusion in concordant topologies. The cloudogram in the top left shows discordance between the wASTRAL (black) and RAxML-NG (gray) topologies. Each genus and subgenus are highlighted in a different color and are represented by a flower image on the right. A scale bar is included for every genus. From top to bottom: *Hillia* subg. *Hillia* (photo **a** [*H. parasitica*] taken by Jason Grant), *Hillia* subg. *Ravnia* (photos **b** [*H. triflora* var. *triflora*] and **c** [*H. longifilamentosa*] taken by Laymon Ball and Barry Hammel, respectively), *Hillia* subg. *Illustres* (photo **d** [*H. illustris*] taken by Laymon Ball), *Hillia* subg. *Tetrandrae* (photo **e** [*H. tetrandra*] taken by Miriam Jiménez and Mariano Gorostiza), *Balmea* (photo **f** from Mejía-Jiménez et al., 2022), and *Cosmibuena* (photo **g** [*C. valerii*] taken by Laymon Ball). Icons to the right of taxon names depict species’ flower shapes.

Three species, *H. macrophylla*, *H. parasitica,* and *H. triflora,* were recovered as non-monophyletic in both the wASTRAL and RAxML-NG phylogenetic analyses. In the case of *H. macrophylla*, one individual collected in Central America was sister to the rest of subgenus *Hillia* in the wASTRAL tree, while the other individuals, all from South America, formed a grade successively sister to *H. wurdackii*, a species with suspected affinity to *H. macrophylla* (Taylor 1994). The individuals of *H. parasitica* form a clade in which two species (*H. killipii* and *H. pumila*) are embedded. Within this clade, two Caribbean *H. parasitica* individuals were sister to one another and the South American representative was sister to *H. killipii*, also from South America.

Most branches had quartet concordance (QC) scores above 0.2 (-0.30–1; x^̅^=0.42), which indicates good support for the species tree topology (Fig. 1; Pease 2018). All branches had a quartet informativeness (QI) score above 0.95 (range: 0.99–1; x^̅^=0.99), suggesting that low information was not an important source of gene tree discordance in our data. The quartet fidelity (QF) score for taxa ranged from 0.56 to 0.91 (x^̅^=0.72). Higher QF scores indicate that when sampled in quartets, taxa tended to produce a topology that was concordant with the species tree; this approach is similar to a “rogue taxon” test (Pease 2018). The quartet differential (QD) score, which measures the skewness in the frequencies of the two discordant quartet topologies, ranged from 0–1 (x^̅^=0.36).

We estimated that Hillieae originated during the early Miocene, 19.1 Ma (95% HPD = 11.1– 26.6; Fig. 2a, b; Supplementary Fig. S2 and Table S4). The crown age of *Cosmibuena* was estimated to be 11.8 Ma (95% HPD = 5.9–18.1). *Balmea* and *Hillia* diverged 16.4 Ma (95% HPD = 9.0–23.7) and the crown age of *Hillia* was estimated to be 10 Ma (95% HPD = 5.3–14.9).

**Figure 2.**
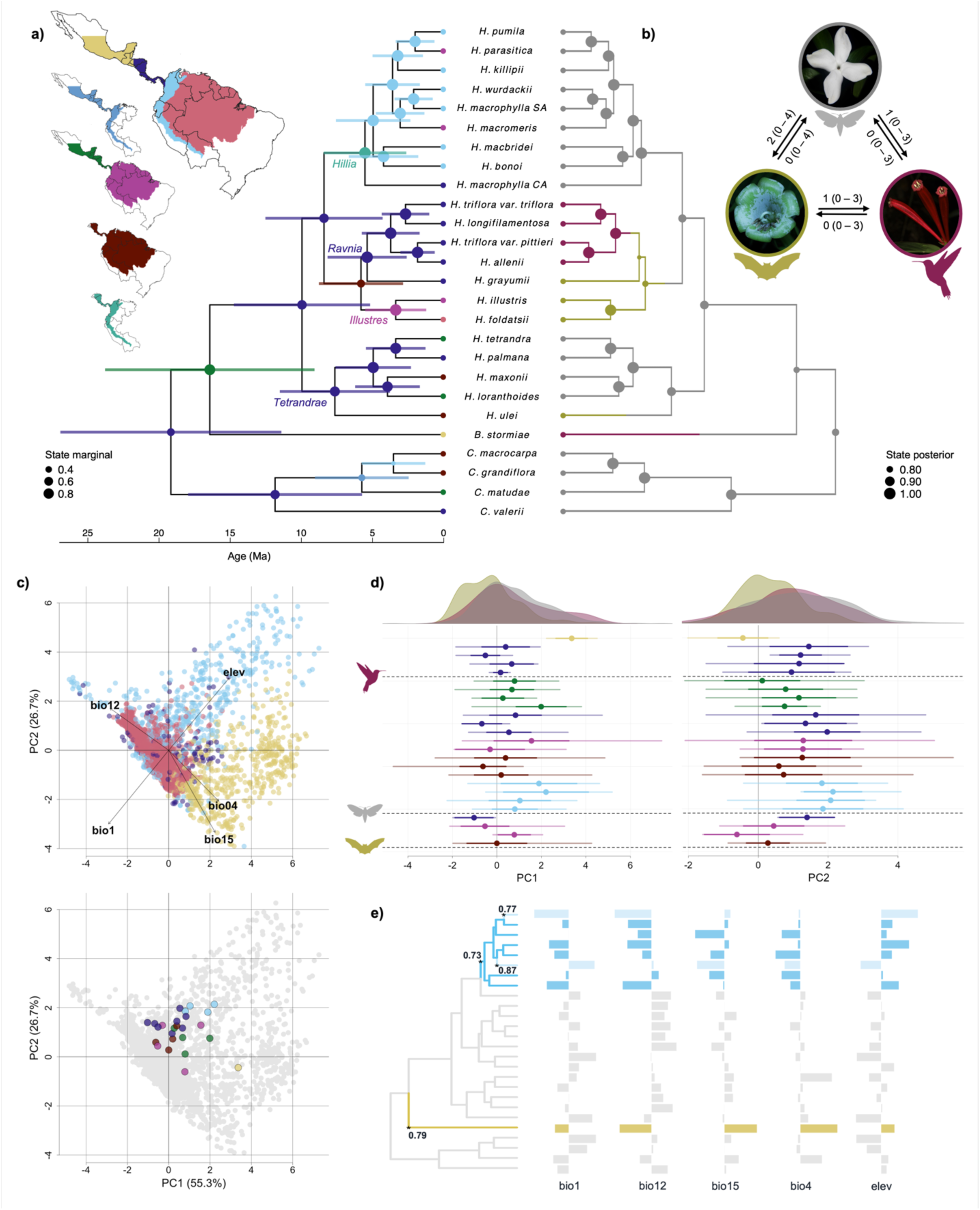
Biogeographic, pollination syndrome, and bioclimatic niche evolution in Hillieae. In **a** and **b**, sizes of the circles at the nodes indicate marginal and posterior probabilities of the ancestral state, respectively. In **a**, the lengths of the bars at the nodes represent the 95% HPD. **a)** Biogeographic modeling (DEC) results. *Hillia* subgenera are labeled. **b)** Pollination syndrome evolution. Plot in the upper right corner shows the number of transitions between syndrome states (with confidence intervals in parentheses). Images of different species are used to exemplify each state (gray = hawkmoth [*H. tetrandra*]; purple = hummingbird [*H. triflora var. triflora*]; green = [*H. illustris*]). **c)** PCA results of bioclimatic variables on the land pixels covering Hillieae’s distribution. On the upper plot, pixels are colored according to geographic region. A random sample of 5,000 pixels (of 455,929) are included for visualization. Variable loadings scaled by a factor of 6 for visibility are also included (bio1 = annual mean temperature; bio4 = temperature seasonality; bio12 = annual precipitation; bio15 = precipitation seasonality; elev = elevation). The lower plot includes species’ averaged bioclimatic niche positions, and points are colored according to region. **d)** Species’ mean niche positions (points), ranges (transparent lines), and standard deviations (solid lines) along each PC axis. Points and lines are colored by geographic region and grouped by pollination syndrome. The density plots above each panel, shaded by pollination syndrome, depict niche variability across individuals for each syndrome along the corresponding PC axis. **e)** Bioclimatic niche shifts detected via *l1ou* analysis. Shifts are colored according to species’ geographic range (*Balmea stormiae* = yellow; Andean *Hillia* subg. *Hillia* = light blue; additional shifts within *Hillia* subg. *Hillia* are light blue and transparent. Bootstrap support values for each shift are shown at the corresponding nodes of the tree.

**Figure 3.**
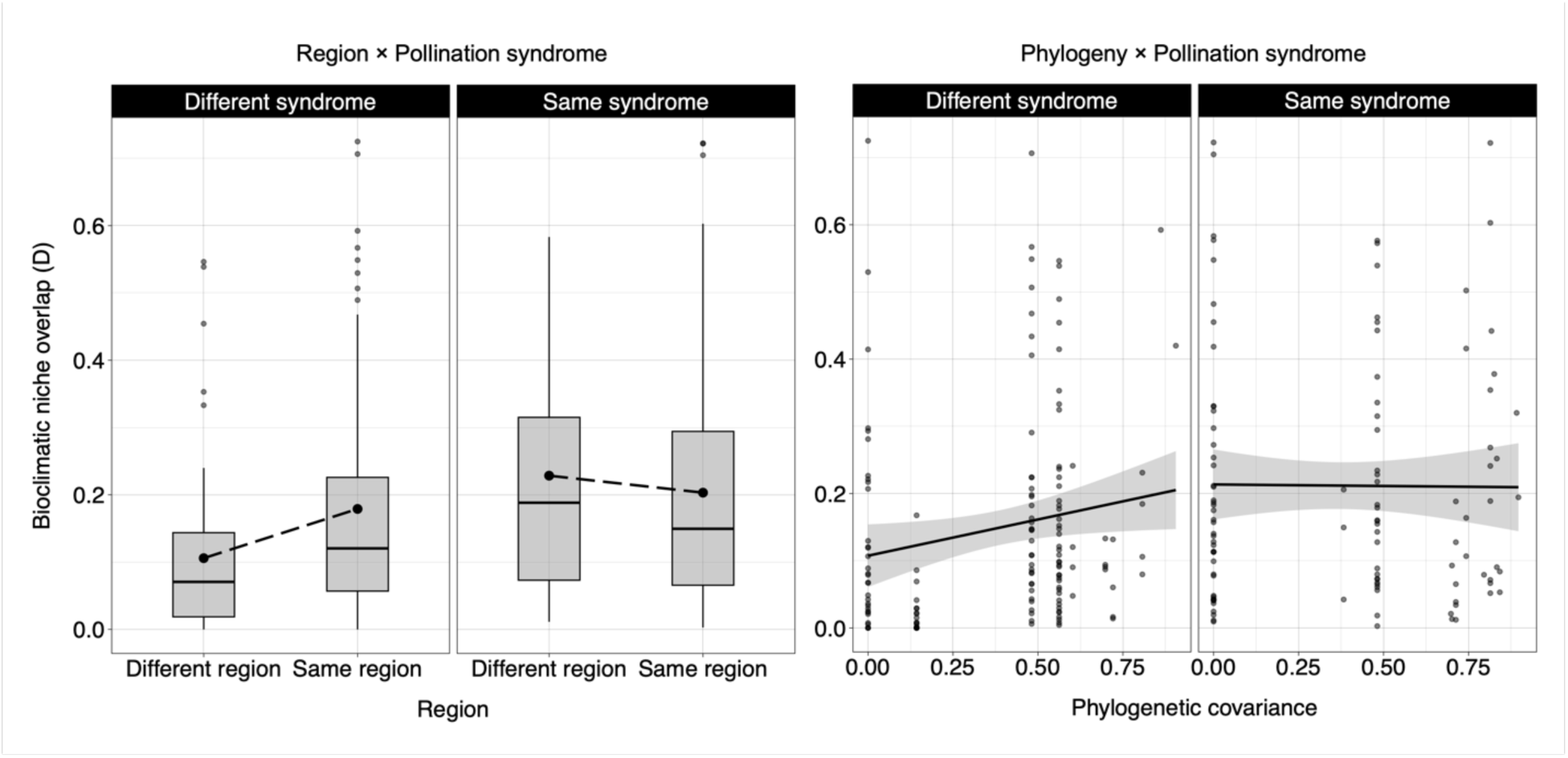
Results from generalized linear model demonstrating additive effects of two interactions on species’ pairwise niche overlap. Left: Interaction between region and pollination syndrome. Species in the same region tend to have greater niche overlap, except when they also share the same pollination syndrome (p=0.001). **Right:** Interaction between phylogenetic covariance and pollination syndrome. Niche overlap generally increases with phylogenetic relatedness, but this effect is weaker when species share the same pollination syndrome (p=0.0002).

### Biogeographic modeling in Hillieae

The DIVA-like (Dispersal–Vicariance Analysis) model was the best-supported based on having the lowest AIC, however, it differed only slightly from the DEC model (ΔAIC < 2). We present results from the DEC. Marginal support for ancestral ranges ranged from 0.34 to 1 (x^̅^=0.84) across all nodes (Fig. 2a; Supplementary Fig. S2 and Table S4). Southern Central America was estimated as the most probable geographic range at the root of Hillieae and its subgenera. Our results indicate that Hillieae have undergone many biogeographic movements within the Neotropics. When *Cosmibuena* split from the rest of Hillieae, the genus continued to diversify primarily within Central America until around 5.72 Ma (95% HPD 2.5–9.1), when the ancestor of the subclade excluding *C. valerii* expanded its range into the Andes (marginal probability = 0.42). A vicariance event likely separated the lineage leading to *C. matudae* (restricted to northern and southern Central America) from the ancestor of *C. macrocarpa* and *C. grandiflora* (restricted to the Andes), and the latter two species independently expanded their ranges back into southern Central America, as well as the Amazon; this is the only instance where Hillieae returned to southern Central America after its dispersal from the region. Prior to the split between the two genera around 16.4 Ma (95% HPD 9.0–23.7), the ancestor of *Hillia* and *Balmea* expanded its range from southern Central America into northern Central America (marginal probability = 0.88). This expansion was followed by a vicariance event that left *Balmea* restricted to northern Central America and *Hillia* to southern Central America. Crown *Hillia* originated 9.9 Ma (95% HPD 5.3–14.9) and primarily diversified within southern Central America. At approximately 5.8 Ma, the ancestor of subgenera *Ravnia* and *Illustres* expanded its range into the Andes and the Amazon (marginal probability = 0.51). A subsequent vicariance event restricted subgenus *Illustres* to the Andes and Amazon, and *Ravnia* to southern Central America. The largest subgenus of *Hillia*, *Hillia* subg. *Hillia*, expanded its range from southern Central America into the Andes 5.5 Ma (marginal probability = 1), after which point the group continued to diversify primarily within the Andes. In total, Hillieae have independently dispersed north from southern Central America into northern Central America at least four times and south into the Andes at least five times; dispersal from the Andes into the Amazon has taken place at least seven times.

### Bioclimatic Niche Evolution

The first two axes of the PCA explained a total of 82% of the climatic variation over Hillieae’s distribution (Fig. 2c). PC1 explained 55.3% of the variation and was positively correlated with temperature seasonality (bio4), precipitation seasonality (bio15), and elevation, and negatively correlated with annual mean temperature (bio1) and annual precipitation (bio12). PC2 explained 26.7% of the variation and was positively correlated with annual precipitation and elevation, and negatively correlated with annual mean temperature, temperature seasonality, and precipitation seasonality (Fig. 2c; Supplementary Table S5; Supplementary Fig. S3). When projected onto the PCA, most species (except *Balmea stormiae*) clustered closely together. Plants with the bat pollination syndrome tend to have lower PC1 and PC2 scores than those with hawkmoth or hummingbird syndromes (Fig. 2d). However, these differences were not statistically significant, regardless of whether phylogeny was accounted for (ANOVA: p>0.05 for both axes; MCMCglmm: pMCMC>0.05 for both axes and all syndrome categories). These results suggest that pollination syndrome alone does not strongly explain variation in species’ realized niches.

Schoener’s index of niche overlap (D) between species ranged from 0 to 0.72 (x^̅^=0.17; Supplementary Table S7). The most complex model (df=9), which includes a three-way interaction between phylogenetic relatedness (i.e., phylogenetic covariance), pollination syndrome, and biogeographic region, had the lowest BIC (Supplementary Table S8). However, this did not drastically differ from the next best-fitting, simpler model (df=7), which includes interactions between pollination syndrome and biogeographic region and between pollination syndrome and phylogenetic relatedness, both additively affecting niche overlap. The difference in BIC between these models was 2.99, indicating positive, but not strong, support for the more complex model (Raftery 1995). Ultimately, we selected the simpler model, which explained 21.3% of the total variance in the data, and had slightly better residual uniformity than the more complex model (Kolmogorov-Smirnov test p-value = 0.68 for the simpler model and 0.47 for the more complex). Individually, closely related species exhibit significantly higher niche overlap than distantly related ones (p=8.20×10⁻^7^); species sharing the same pollination syndrome have significantly higher niche overlap than those with different syndromes (p=9.49×10⁻^11^); and species occurring in the same region show significantly higher niche overlap than those in different regions (p=4.64×10⁻^5^). When species share the same pollination syndrome and occur in the same region, the interaction between these variables negatively affects niche overlap (p=0.001). Similarly, when species are closely related and share the same pollination syndrome, the interaction between the variables results in a combined negative effect (p=0.0002).

We detected four evolutionary shifts in bioclimatic niche with *l1ou*: one shift was in *Balmea stormiae*, and the other three shifts were in the mainly Andean subclade of *Hillia* subg. *Hillia* (Fig. 2e; see Supplementary Fig. S4 and Supplementary Table S6 for species-specific distributions of bioclimatic variables).

### Pollination Syndrome Evolution

Hawkmoth pollination syndrome is inferred as the ancestral character state for Hillieae with moderate posterior probability (0.82). There have been two (95% CI: 0–4) shifts from hawkmoth to bat, one shift from bat to hummingbird (95% CI: 0–3), and one (95% CI: 0–3) shift from hawkmoth to hummingbird pollination syndromes. Posterior support for ancestral syndrome states ranged from 0.78–1 (x^̅^ =0.94) across all nodes (Fig. 2b; Supplementary Fig. S2 and Table S4). All repeated shifts were likely the result of convergence, as we did not find evidence for hemiplasy in our dataset— only two gene trees showed monophyly with respect to the bat syndrome (i.e., *Hillia ulei* formed a monophyletic clade with bat-syndrome *Hillia*. subg. *Ravnia*). No gene trees showed monophyly with respect to the hummingbird syndrome (i.e., within the gene trees, *Balmea* never formed a monophyletic clade with hummingbird-syndrome *Hillia* subg. *Ravnia*).

## Discussion

### Systematics and Taxonomy

We infer the first densely sampled, well supported phylogeny for tribe Hillieae using target enrichment sequencing and a Rubiaceae-specific probe-set designed to target 2270 loci. We found that current taxonomy (i.e., generic and species descriptions) was consistent with phylogeny with a few exceptions. *Cosmibuena* and *Hillia* are recovered as monophyletic, with *Cosmibuena* sister to both *Hillia* and monotypic *Balmea*. This result corroborates previous molecular studies which have only included 1-3 species from each genus (Robbrecht and Manen 2006; Manns and Bremer 2010; Manns et al. 2012; Paudyal et al. 2014). In our phylogenetic analysis we recovered four subgenera slightly modified from Taylor 1994, which we hereafter refer to as: *Hillia*, *Illustres*, *Ravnia*, and *Tetrandrae*.

Multiple species were inferred as nonmonophyletic. The non-monophyly of both *H. macrophylla* and *H. parasitica* indicates that disjunct lineages in those species may represent distinct species that have become reproductively isolated without notable morphological divergence. In both phylogenetic analyses, the two *Hillia triflora* varieties never formed a clade; our results show *Hillia triflora* var. *pittieri* as sister to *H. allenii* with strong support and *H. triflora* var. *triflora* as sister to *H. longifilamentosa*, but with weak support (the placement of *H. longifilamentosa* within this clade is highly uncertain). This suggests that these two varieties may, in fact, represent two distinct species. Alternatively, incomplete lineage sorting (ILS) or gene flow in areas of sympatry can also result in the recovery of species as paraphyletic. Our QuartetSampling results suggest that these are both plausible scenarios contributing to gene-tree discordance and paraphyly, particularly for *H. parasitica*. All three *H. parasitica* individuals (in addition to the rest of the taxa in its subgenus, *Hillia*) have low quartet fidelity scores which could indicate erroneous taxa placement, or that lineage-specific processes are distorting the phylogeny (Pease et al. 2018). Additionally, many of the branches within this clade have low quartet concordance (<0.2), and quartet differential (QD) scores are highly variable, suggesting that a combination of ILS and introgression could be causing discordance and non-monophyly of species. ∼28% of quartets sampled for the branch leading to *H. killipii* and one *H. parasitica* support an alternative topology where all *H. parasitica* and *H. pumila* are monophyletic. The low QD score (0.29) at this branch is also consistent with introgression causing discordance.

However, it should be noted that low QD can also reflect discordance due to other underlying biological processes such as heterogeneity in evolutionary rates or base compositions (Pease et al. 2018). Ongoing work at the population level for these species will help unravel geographical patterns of genetic variability and inform species delimitation to support taxonomic revision.

### Biogeographic History and Bioclimatic Niche Evolution

Since their origin in southern Central America about 19 Ma (95% HPD 11.1–26.6), Hillieae have had a dynamic biogeographic history characterized by frequent dispersal between Central and South America. This is expected, given that these species are centered in geologically and ecologically dynamic regions: southern Central America, which has some of the youngest ecosystems in the Neotropics, and the Northern Andes, which is the fastest uplifting portion of the continent-spanning mountain range (Gregory-Wodzicki 2000; Garzione et al. 2008; Pérez-Escobar et al. 2022). Additionally, ten of the 26 species sampled in the phylogeny have ranges spanning at least two major regions, with four extending across the Andes mountains—a well-known barrier to dispersal for many other plant and animal groups. Hillieae’s wind-adapted seeds (i.e., small, papery, and lightweight) and epiphytic habit, which allows seed release from elevated heights, have likely facilitated its dispersibility, a pattern observed in other epiphytic groups (Kessler 2002; Mountier et al. 2018)

Hillieae diversification was contemporaneous with three major geological events: the bridging of the Isthmus of Panama, mountain uplift in southern Central America, and the final uplift of the northernmost Andes. Coupled with Hillieae’s high dispersibility, these events would have likely played a significant role in structuring evolutionary relationships. Based on our biogeographic modeling results, we estimate that three independent dispersal events across the Panamanian Isthmus into South America occurred ∼6 Ma, involving the ancestors of *Hillia* subg. *Hillia*, *Ravnia* + *Illustres*, and *Cosmibuena* (excluding *C. valerii*). Additionally, four extant taxa (*H. maxonii*, *H. ulei*, *C. macrocarpa*, and *C. grandiflora*) expanded their ranges across the Isthmus. The fact that multiple lineages dispersed across the Isthmus at approximately the same time supports a slightly earlier timing of Isthmus closure (at least before 5 Ma), and challenges the traditional view of a later closure around 3.5 Ma (Bacon et al. 2015). However, because these are wind-dispersed epiphytes, we cannot rule out the alternative explanation that long-distance dispersal over a discontinuous land bridge facilitated their expansion.

Showing an important role of the landscape on diversification within Hillieae, the predominantly Andean clade, *Hillia* subg. *Hillia*, experienced rapid radiation relative to the rest of the tribe since its origin *ca*. 5.5 ma. This is inferred from the short internodes, lower support, and the relatively high levels of gene tree discordance observed within the clade, which is typical of rapidly radiating Andean-centered groups (Lagomarsino et al. 2022; Tribble et al. 2024).

Differences in bioclimatic niche across Hillieae reflect the highly variable and dynamic nature of the Neotropical landscape, as well as the biology of the group. Hillieae are distributed throughout the Neotropics, though most species are constrained to a central region of the total available niche space, preferring mid-elevation wet forests with low seasonality in both precipitation and temperature. The prevalence of bioclimatic niche conservatism among most species may be indicative of constraints imposed by their epiphytic habit. Because many epiphytes capture rainwater either directly from rainfall or from the surfaces on which they grow rather than from the soil, their water supplies tend to be limited, and depending on the group, water stress can be even initiated by rainless periods of just a few hours in some epiphytic groups (Zotz and Hietz 2001; Spicer and Woods 2022). This limitation in water availability may constrain the seasonal regimes within which most Hillieae are able to establish themselves, with less seasonal (i.e., more consistent rainfall) regions being more favorable (Kreft et al. 2004). Consistent with this, species are more variable in their total annual precipitation and elevation preferences than they are in seasonality, as indicated by our PCA. We detect two major evolutionary shifts in bioclimatic niche that happen contemporaneously with shifts in biogeography. One shift occurs in a subclade of *Hillia* subg. *Hillia* concurrently with its dispersal into the Andes and indicates movement into to cooler, drier, high elevation habitats with low seasonality. The other occurs in *Balmea stormiae*, and indicates an adaptation to higher elevation, cooler, drier, and more seasonal habitats. This shift is contemporaneous with its dispersal to northern Central America— the northernmost distribution in Hillieae. *Balmea stormiae* is also the only Hillieae species that is variably terrestrial or epiphytic (Taylor 1994; Lorence and Taylor 2012; Mejía-Jiménez and Montero-Castro 2022), which likely minimizes water stress in these relatively seasonable environments.

### Pollination Syndrome Evolution

Using color as a proxy, we found repeated shifts between bat, hawkmoth, and hummingbird pollination syndromes in the group. The hummingbird and bat pollination syndromes evolved from hawkmoth pollinated ancestors (once and twice, respectively), and there has been a single shift to the hummingbird pollination syndrome from the bat syndrome; no reversals to the hawkmoth syndrome have occurred. Certain ecological, morphological, physiological, or phenological traits can facilitate or inhibit shifts between pollination modes (Bradshaw and Schemske 2003; Anderson and Johnson 2009; Vereecken 2009; Whittall and Carlson 2009; van der Niet and Johnson 2012; Johnson et al. 2017; Givnish et al. 2020). For example, tubular corollas are components of bat, hawkmoth, and hummingbird pollination, and nocturnal flowering with scent production are components of both hawkmoth and bat pollination. It is likely that aspects of the hawkmoth pollination syndrome facilitated shifts from hawkmoth to other pollination syndromes in Hillieae.

While floral morphology tends to align with primary floral visitors, deviations from expectations can occur when species are pollinated by multiple functional groups (Stebbins 1970; Fenster et al. 2004; Rosas-Guerrero et al. 2014; Ashworth et al. 2015). Among the species sampled in the current study, two species (*Balmea stormiae* and *Hillia allenii*) have trait combinations that make it difficult to exclusively assign them to a single syndrome: *Balmea stormiae* has tubular red to purple flowers consistent with hummingbird pollination, and is known to produce sweet-smelling odors at night, which is typical for hawkmoth pollination; and *H. allenii* has a wide funnelform flower that is typical of bat pollination, and a highly visible salmon-colored flower that is consistent with hummingbird pollination, though these are sometimes partially pale green-yellow (Martínez 1942; Mejía-Jiménez and Montero-Castro 2022; observations from herbarium specimens). Both species were coded as hummingbird pollinated based on their primary floral color, though we suspect that their combined traits could either reflect recent transitions (hawkmoth to hummingbird for *B. stormiae* and hummingbird to bat for *H. allenii*) between pollination modes, or mixed pollination modes. These species underscore the nonbinary nature of syndromes (Brightly et al. 2024) and field studies to assess whether their trait combinations can be attributed to selection by multiple pollinators in *B. stormiae* and *H. allenii* is an ongoing and particularly promising line of future research.

Hemiplasy has been implicated in apparent trait convergence in some groups (Hibbins et al. 2020), including in the context of pollination syndrome evolution (Stankowski and Streisfeld 2015). However, we find no evidence for hemiplasy in the evolution of pollination syndromes in Hillieae, at least not within the set of Rubiaceae2270x loci included in this study. The frequency of pollination syndrome shifts in Hillieae suggests that modifying floral traits may not be particularly challenging for this group, potentially due to relatively flexible developmental pathways. Hillieae’s small clade size and the repeated evolution of pollination syndromes within it makes it a promising system for understanding the genetic basis of pollination syndromes when better genomic resources are available.

### Interplay Between Abiotic and Biotic Factors in Hillieae Evolution

Geographical patterns of pollination syndrome diversity point to the interplay between biogeographic history, climatic preferences and pollination in shaping Hillieae diversification. Hillieae’s species and floral diversity is concentrated in southern Central America, where the group originated. This is the only region where all three pollination syndromes are represented (including a mixed-syndrome species, *Hillia allenii* from subgenus *Ravnia*) and multiple shifts between syndromes have occurred. In contrast, species that diversified within the Andes (i.e., *Hillia* subg. *Hillia*) display the ancestral hawkmoth syndrome. Curiously, no hummingbird-syndrome Hillieae occur in the Andes (i.e., all Andean Hillieae are either salverform and white, or funnelform and green) even though hummingbird pollination is common in this region, and the proportion of hummingbird pollinated taxa is expected to increase with elevation (Cruden 1972; Dellinger et al. 2023; Barreto et al. 2024). Notably, species in *Hillia* subg. *Hillia* occur at markedly higher elevations than Central American hummingbird-syndrome Hillieae. Despite originating around the same time (approximately 5.8 and 5.5 Ma, respectively), the clade comprising *Hillia* subgenera *Illustres* and *Ravnia* has experienced more pollination syndrome shifts compared to its sister clade, subgenus *Hillia*, even though the latter has higher extant species richness. The disparity in pollination syndrome diversity between southern Central America and the Andes likely reflects a combination of evolutionary contingency and regional differences in the influence of other abiotic (e.g., habitat heterogeneity, topographic barriers) and biotic (e.g., species competition) factors, potentially mediated by geological differences between the two regions.

Compared to Central American mountain ranges, the Andes are characterized by much higher and more extensive topography. These mountains, which are the longest continental mountain range in the world, stretch across the western edge of South America and have an average height of 4000 m. The extreme topography of the Andes lends itself to high habitat and local climate variation (Huddart and Stott 2020), which have been key abiotic drivers of rapid diversification events in this region (Linder 2008; Luebert et al. 2011; Jabaily and Sytsma 2012; Givnish et al. 2014, 2015; Sánchez-Baracaldo and Thomas 2014; Lagomarsino et al. 2016; Pouchon et al. 2018). Throughout the landscape, high peaks with dry and cold habitats are juxtaposed with warmer, wet or dry river valleys, which can serve as geological barriers separating populations and species (Hazzi et al. 2018). While this geological diversity is also present in southern Central America, it exists at a much smaller scale in comparison. Some studies have found a positive relationship between topographic complexity and intraspecific genetic variation (Guarnizo and Cannatella 2013; García-Rodríguez et al. 2021; Dellinger et al. 2022), which can lead to speciation. The environmental heterogeneity of the Andes is reflected in the bioclimatic niche diversity of the predominantly Andean clade, *Hillia* subg. *Hillia*— while these species do not show major differentiation on the climatic PCA, the *l1ou* analysis based on separate bioclimatic variables shows that multiple niche shifts have occurred within this group.

Given their geological differences, the Andes and southern Central America may promote coexistence among Hillieae species through different mechanisms. Our best-fitting generalized linear model (GLM) indicates that same-syndrome species pairs tend to exhibit greater climatic niche overlap, except when species co-occur in the same region or are closely related. This pattern could reflect niche partitioning driven by competition between species in proximity for pollinator resources, or a mechanism of reproductive reinforcement to maintain species boundaries. For Andean *Hillia* subg. *Hillia*, the heterogeneous nature of the Andes has likely provided abundant opportunity for allopatry and niche differentiation, allowing for diversification without major divergences in pollinator niche. In contrast, floral variation and pollination syndrome diversity in Southern Central American Hillieae likely reduces direct competition between species for pollinators and allows for coexistence over a smaller and relatively more homogenous topography that is also highly species rich. Variation in floral morphology has been observed to reduce competition among co-occurring taxa in multiple systems (Rodríguez-Gironés and Santamaría 2007; Armbruster and Muchhala 2009; Muchhala et al. 2014; Johnson et al. 2017).

## Conclusions

We present the first robust molecular phylogeny for tribe Hillieae, shedding light on a dark branch of the Tree of Life. Using this new phylogenetic framework, we characterized the evolution of pollination syndrome, biogeography, and bioclimatic niche within the tribe. Our results show that while bioclimatic niche has been largely conserved, with just four major evolutionary shifts (two in distant lineages *Balmea* and *Hillia* subg. *Hillia*, and another two within *Hillia* subg. *Hillia*), dispersal across the Neotropics has been dynamic, and multiple independent shifts between bat, hawkmoth, and hummingbird pollination syndromes have occurred. Notably, some of these shifts have occurred concurrently, highlighting the link between abiotic and biotic factors in driving Hillieae evolution. When species share pollination syndromes, climatic niche differentiation may facilitate coexistence of species either when they occur in the same biogeographic region or they are closely related. Neotropical topography has likely played a central role in shaping the ecological contexts of diversification, indicated by regional disparities in pollination syndrome diversity. We hope that this work will inform future evolutionary analyses in the tribe and broader Rubiaceae while offering insight into broader patterns of flowering plant diversification in the Neotropics.

**Dryad link for reviewers:** https://datadryad.org/stash/share/8e05bFmAvnKk_W29-IO7fw1oM7xFbWdEMMAlj5E119E

## Supplementary Material

Data available on Dryad (https://doi.org/10.5061/dryad.4mw6m90k6).

**Supplementary Figure S1**. Concatenated tree inferred with RAxML-NG. Felsenstein’s bootstrap proportions (FBP) less than 100 are plotted above branches.

**Supplementary Figure S2.** Phylogeny labeled with node numbers for reference in

**Supplementary Figure S3.** Geographic distribution of each bioclimatic variable included in niche analyses. Maps illustrate spatial coverage and variation across the study region.

**Supplementary Figure S4.** Boxplots showing species-specific distributions of WorldClim bioclimatic variables. Boxplot colors correspond to colors in the maps to the right. **Bio1** = annual mean temperature; **Bio4** = temperature seasonality; **Bio12** = annual precipitation; **Bio15** = precipitation seasonality.

**Supplementary Table S1. Accession information for sampled Hillieae.** Grayed rows indicate specimens that failed sequencing. For institutions: LSU= Louisiana State University and MO= Missouri Botanic Garden. Samples are a part of NCBI BioProject PRJNA970862.

**Supplementary Table S2. HybPiper summary statistics.** Grayed rows indicate samples with assembled contigs for less than 10% of loci.

**Supplementary Table S3.** AMAS alignment statistics for the final 1375 processed loci. Average values for each statistic, followed by the minimum and maximum values, are shown.

**Supplementary Table S4. Results for divergence dating analysis and ancestral state reconstructions of biogeography and pollination syndrome.** Clade ages (estimated via RevBayes) in Millions of Years; 95% Highest Posterior Density (HPD) Intervals; geographic ranges inferred with BioGeoBEARS at nodes (A = Andes, B = southern Central America, C = northern Central America, and D = Amazon) and their marginal probabilities; pollination syndromes inferred with RevBayes at nodes and their posterior probabilities. See Supplementary Fig. S2 for wASTRAL species tree topology with node numbers.

**Supplementary Table S5. Relative contributions of each variable to the bioclimatic PCA axes (i.e., variable loadings).** bio1 = annual mean temperature; bio4 = temperature seasonality; bio12 = annual precipitation; bio15 = precipitation seasonality; elev = elevation.

**Supplementary Table S6. Summary statistics for each species and WorldClim bioclimatic variable included in the niche analyses.** (A = Andes, B = southern Central America, C = northern Central America, and D = Amazon). Bolded rows indicate species with less than 5 occurrence records. Species names with an asterisk (*) were not sampled in the phylogeny.

**Supplementary Table S7. Results of pairwise niche overlap analysis.** Table includes the species pairs, whether they share the same pollination syndrome and geographic range, and their Schoener’s index (D) of niche overlap.

**Supplementary Table S8. Statistics for 18 generalized linear models of pairwise niche overlap between species: df** = degrees of freedom; **ΔBIC** = the difference in Bayesian Information Criterion (BIC) between the model and the model with the minimum BIC; **phylogeny** = the coefficient for the relationship between phylogenetic covariance and species pairwise niche overlap; **syndrome** = the coefficient for when species share the same pollination syndrome; **region** = the coefficient for when species occur in the same region; **P × S** = the coefficient for the interaction between phylogeny and syndrome; **P × R** = the coefficient for the interaction between phylogeny and region; **S × R** = the coefficient for the interaction between syndrome and region; **P × S × R** = the coefficient for the interaction among phylogeny, syndrome, and region; and **R²** = Ferrari’s pseudo R² for beta-regression models. Significance levels for p-values are as follows: *** (p ≤ 0.001), ** (0.001 < p ≤ 0.01), * (0.01 < p ≤ 0.1), and no asterisk (p > 0.1). Cells with grayed “na” indicate that the variable or interaction was not included in that particular model.

