## Supplementary Figures for "Phylogenomics and macroevolution of a florally diverse Neotropical plant clade (Hillieae)"

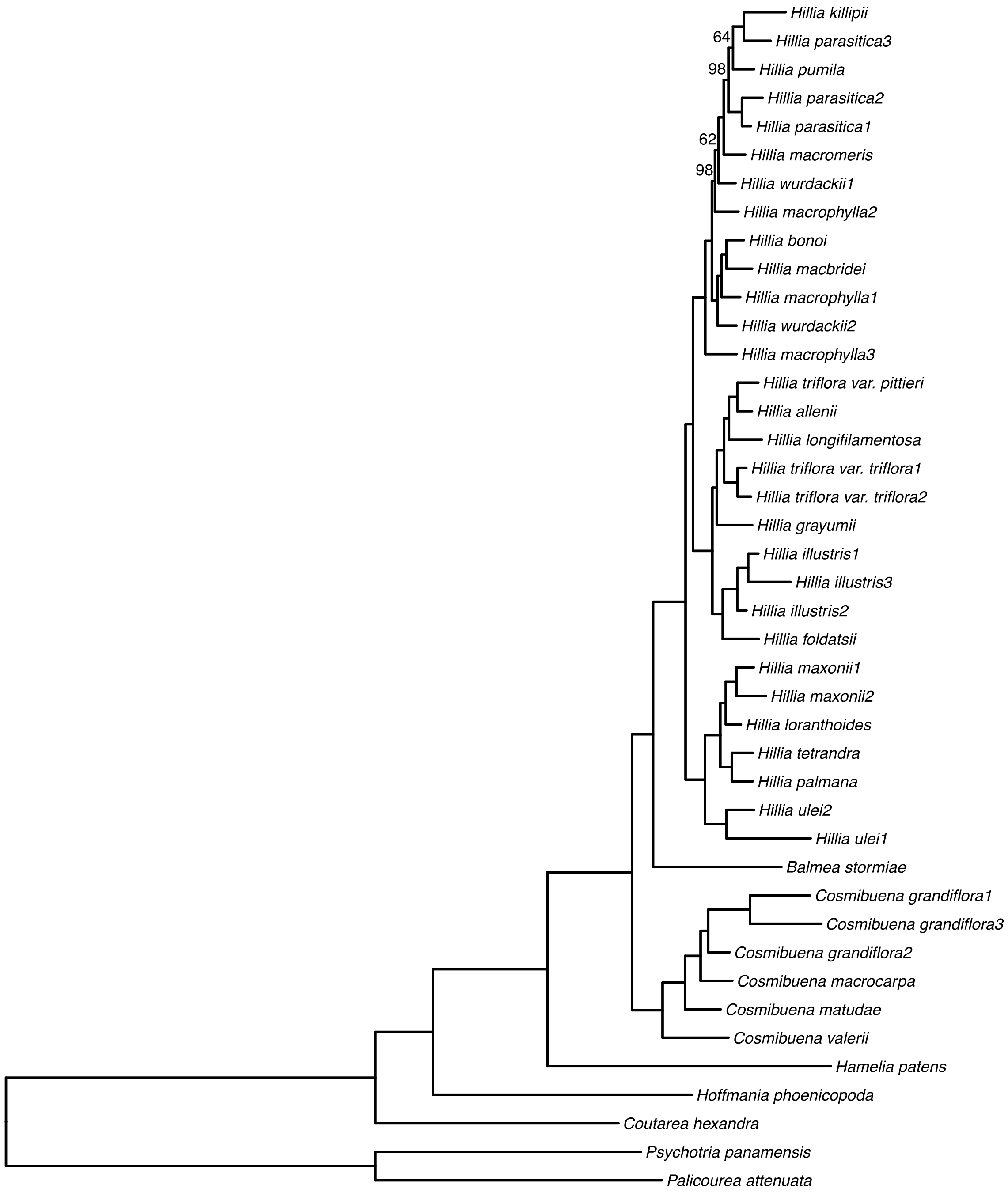


**Supplementary Figure S1.** Concatenated tree inferred with RAxML-NG. Felsenstein's bootstrap proportions (FBP) less than 100 are plotted above branches.

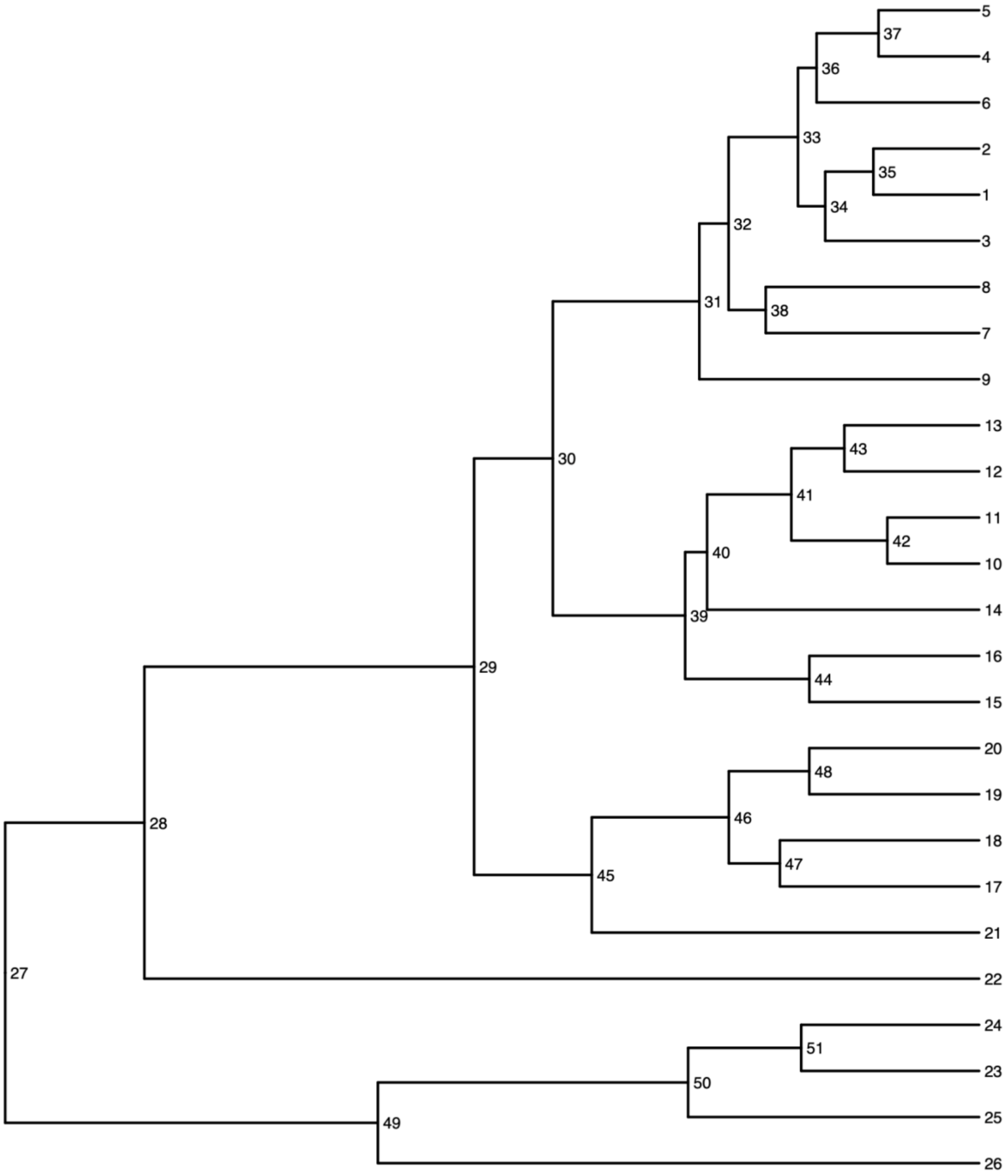


**Supplementary Figure S2.** Phylogeny labeled with node numbers for reference in Supplementary Table S4.

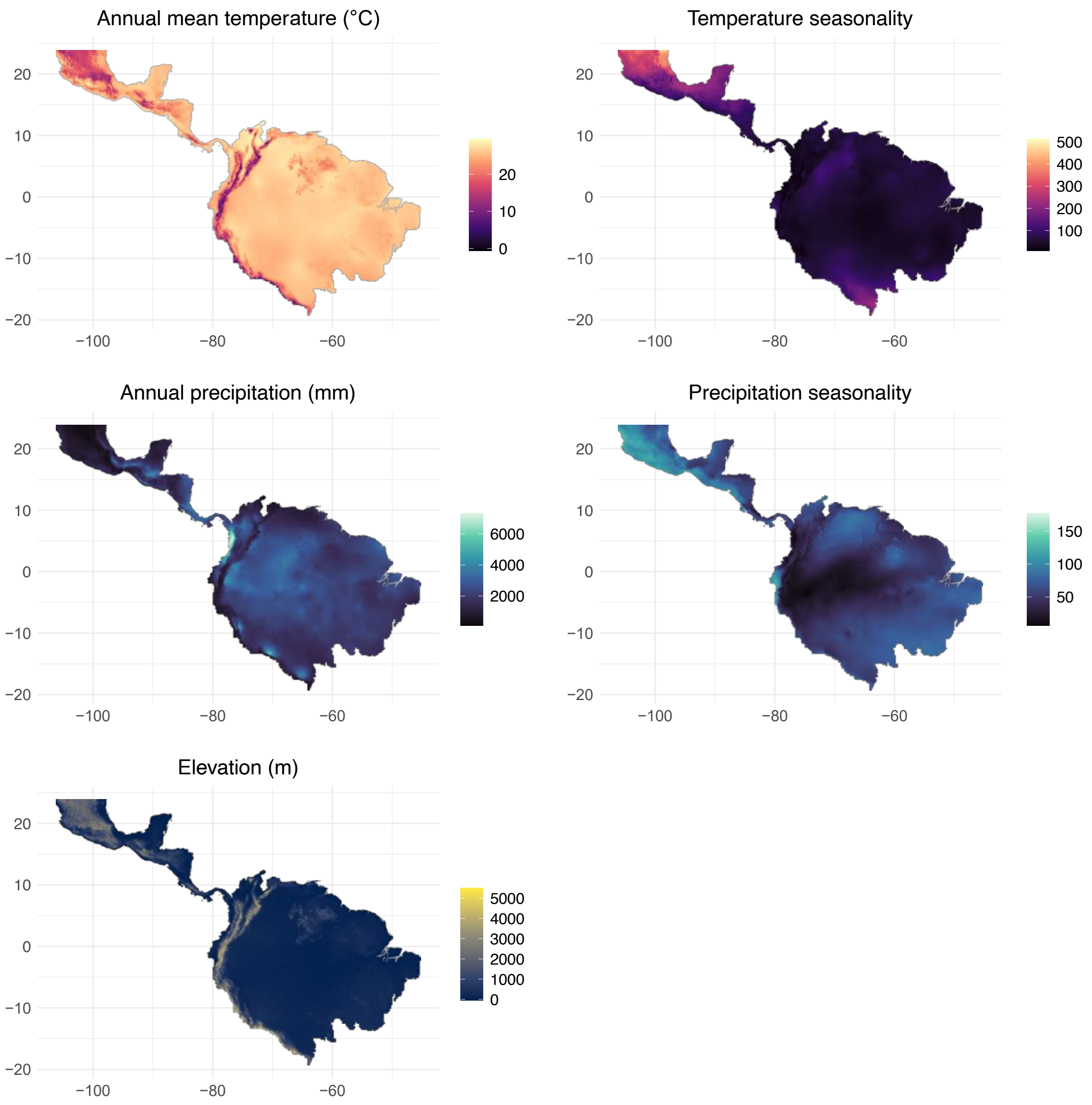


**Supplementary Figure S3.** Geographic distribution of each bioclimatic variable included in niche analyses. Maps illustrate spatial coverage and variation across the study region.

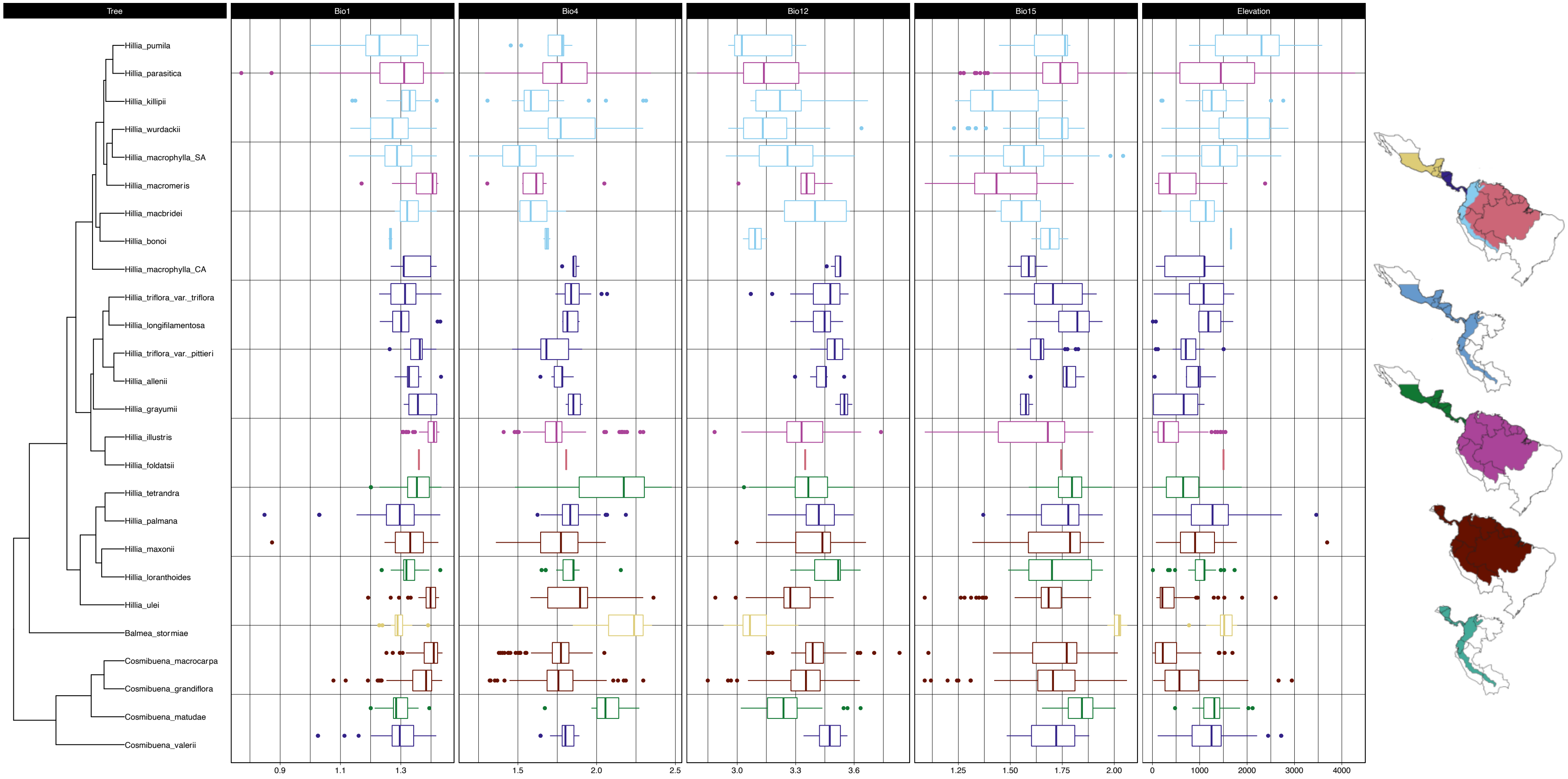


**Supplementary Figure S4.** Boxplots showing species-specific distributions of WorldClim bioclimatic variables. Boxplot colors correspond to colors in the maps to the right. **Bio1** = annual mean temperature; **Bio4** = temperature seasonality; **Bio12** = annual precipitation; **Bio15** = precipitation seasonality.
